# Functional genomic analysis of adult and pediatric brain tumor isolates

**DOI:** 10.1101/2023.01.05.522885

**Authors:** Pia Hoellerbauer, Matt C. Biery, Sonali Arora, Yiyun Rao, Emily J. Girard, Kelly Mitchell, Pratiksha Dighe, Megan Kufeld, Daniel A. Kuppers, Jacob A. Herman, Eric C. Holland, Liliana Soroceanu, Nicholas A. Vitanza, James M. Olson, Justin R. Pritchard, Patrick J. Paddison

**Affiliations:** Human Biology Division, Fred Hutchinson Cancer Center, Seattle, WA USA; Molecular and Cellular Biology Program, University of Washington, Seattle, WA USA; Clinical Research Division, Fred Hutchinson Cancer Center, Seattle, WA USA; Ben Towne Center for Childhood Cancer Research, Seattle Children’s Research Institute, Seattle, WA, USA; Huck Institute for the Life Sciences, Pennsylvania State University, State College, PA, USA; California Pacific Medical Center Research Institute, San Francisco, CA 94107, USA

**Keywords:** adult brain tumor, pediatric brain tumor, functional genomic screens, CRISPR-Cas9, networks, atypical teratoid/rhabdoid tumors, diffuse intrinsic pontine glioma, ependymoma, glioblastoma, medulloblastoma

## Abstract

**Background:** Adult and pediatric tumors display stark differences in their mutation spectra and chromosome alterations. Here, we attempted to identify common and unique gene dependencies and their associated biomarkers among adult and pediatric tumor isolates using functional genetic lethal screens and computational modeling.

**Methods:** We performed CRISRP-Cas9 lethality screens in two adult glioblastoma (GBM) tumor isolates and five pediatric brain tumor isolates representing atypical teratoid rhabdoid tumors (ATRT), diffuse intrinsic pontine glioma, GBM, and medulloblastoma. We then integrated the screen results with machine learning-based gene-dependency models generated from data from >900 cancer cell lines.

**Results:** We found that >50% of candidate dependencies of 280 identified were shared between adult GBM tumors and individual pediatric tumor isolates. 68% of screen hits were found as nodes in our network models, along with shared and tumor-specific predictors of gene dependencies. We investigated network predictors associated with ADAR, EFR3A, FGFR1 (pediatric-specific), and SMARCC2 (ATRT-specific) gene dependency among our tumor isolates.

**Conclusions:** The results suggest that, despite harboring disparate genomic signatures, adult and pediatric tumor isolates share a preponderance of genetic dependences. Further, combining data from primary brain tumor lethality screens with large cancer cell line datasets produced valuable insights into biomarkers of gene dependency, even for rare cancers.

**Importance of the Study:** Our results demonstrate that large cancer cell lines data sets can be computationally mined to identify known and novel gene dependency relationships in adult and pediatric human brain tumor isolates. Gene dependency networks and lethality screen results represent a key resource for neuro-oncology and cancer research communities. We also highlight some of the challenges and limitations of this approach.

## Introduction

Finding effective treatments for adult and pediatric brain tumors remains a daunting challenge. For the last three decades, while significant progress has been made for treatment of other pediatric cancers, like leukemia, effective treatments for brain tumors have failed to materialize. Standard of care (SOC) for gliomas (e.g., glioblastoma (GBM)) arising in the cerebral cortex requires patients to undergo surgical resection and chemoradiotherapy, resulting in lengthy recoveries and often significant side effects [1]. Even with SOC, median survival for GBM patients overall ranges from 14-17 months, with rare exceptions of long-term survival [2, 3].

In children, brain tumors are now the leading cause of cancer-related deaths, responsible for 30% of such deaths [4, 5], with gliomas comprising ~60% of cases [6]. The two year survival rate for pediatric gliomas ranges from 30% for tumors arising in the cerebral cortex to <10% for diffuse intrinsic pontine gliomas (DIPGs)[7]. Other pediatric brain tumors include embryonal tumors such as medulloblastoma (MB) and atypical teratoid/rhabdoid tumors (ATRT) and ependymoma (EPN) [8–12].

Recent efforts to characterize molecular features and candidate disease drivers have revealed stark differences in the mutation spectra and chromosome alterations observed adult and pediatric brain tumors (e.g., EGFR and PTEN) [13–21], even where tumors are histologically indistinguishable. Thus, understanding of similarities and differences in functional genetic dependencies between adult and pediatric tumors may be one key to identifying appropriate therapeutic approaches for these respective tumors [22].

As an alternative approach, candidate therapeutic targets have been sought via functional genomic screens to identify synthetic lethal relationships, i.e., novel gene dependencies driven by cancer-specific alterations [23, 24](reviewed in [25]). These approaches have revealed several types of synthetic lethal interactions, including: loss of paralogs (e.g., ARID1A/ARID1B, SMARCA2/SMARCA4)[26, 27], collateral lethality due to the co-deletion of genes near tumor suppressor loci (e.g., PRMT5 and MTAP-CDKN2A) [28, 29], pathway lethality (e.g., PTEN/PIK3CB) [30], and vulnerabilities associated with molecular features (e.g., WRN/microsatellite instability [31]).

One lingering question from these studies is the degree to which serum-derived cancer cell lines reproduce the genomic, molecular, and phenotypic features of their original patient cancers. Standard *in vitro* growth conditions trigger profound transcriptional, epigenetic, and DNA alterations changes (e.g., [32, 33]). By contrast, for GBM, for example, serum-free culture methods that resemble a stem cell niche allow retention of many of the properties associated with patient tumor isolate, including stem-like cell states [32, 34–36].

For brain tumor isolates, serum-free culture methods that resemble a stem cell niche allow retention of many of the properties associated with patient tumor isolate, including stem-like cell states [32, 34–36]. Performing screens in brain tumor isolates in these culture conditions has revealed multiple novel gene dependencies for GBM cells [37–42]. However, the scope of screens in primary cells is limited by more costly and elaborate culture conditions.

To overcome these limitations, here, we combined the results of CRISPR-Cas9 lethality screen results from adult and pediatric brain tumor isolates with network models generated from cancer cell line functional genomic data [43]. The results reveal that these networks can be leveraged to provide insight into genetic dependencies in brain tumor patient isolates, even for rare brain tumors.

## Results

### CRISPR-Cas9 screens in primary adult and pediatric brain tumor isolates

We previously performed genome-scale CRISPR-Cas9 screens in two adult human GSCs (GSC-0131-mesenchymal and GSC-0827-proneural) and two normal diploid NSC isolates (CB660 and U5) to identify genes differentially required for GSC outgrowth isolates [41]. We also performed the same screen using pediatric GSC-1502-mesenchymal cells (unpublished). Using sgRNA-seq to compare pre/post outgrown populations and GSCs versus NSCs, we identified hundreds of candidate GBM-lethal genes, validating a small portion, including the Wee1-like kinase *PKMYT1* [41]. In this study, we created a comprehensive retest library from this data containing sgRNAs targeting: 208 candidate lethal targets from GSC-0131 cells, 763 from GSC-0827s, and 192 from GSC-1502s. Hits were selected based on a relaxed scoring criteria (≥1 sgRNA at logFC<-1 (FDR<.05)) and whether they failed to score in NSCs. In addition, the retest pool contained gene hits from a human embryonic stem cell screen [44] and biological controls (44 genes) and non-targeting controls (110 sgRNAs). In total this library contained 6591 sgRNAs targeting 1079 genes (**Table S1**).

We then screen this pool using the same outgrowth lethality screening approach in adult GBM stem-like isolates and pediatric brain tumor isolates including atypical teratoid/rhabdoid tumors (ATRT), diffuse intrinsic pontine glioma (DIPG), and medulloblastoma (MED). These included GSC-0131, GSC-0827, and GSC-1502 cells (mentioned above), ATRT-310 (SHH subgroup; *SMARCB1* (focal loss)), ATRT-311 (SHH subgroup; *SMARCB1* (stop gain mutation)), DIPG-4E (*ACVR1, H3.1 K27M*), and MED-411 (group 3; *MYC* amplification) [45] (**Figure 1A**). These isolates were chosen because they grow robustly *in vitro* making them amenable for screens. A representative example of the screen results with positive and negatively scoring sgRNAs is shown in **Supplementary Figure S1A**.

**Figure 1:**
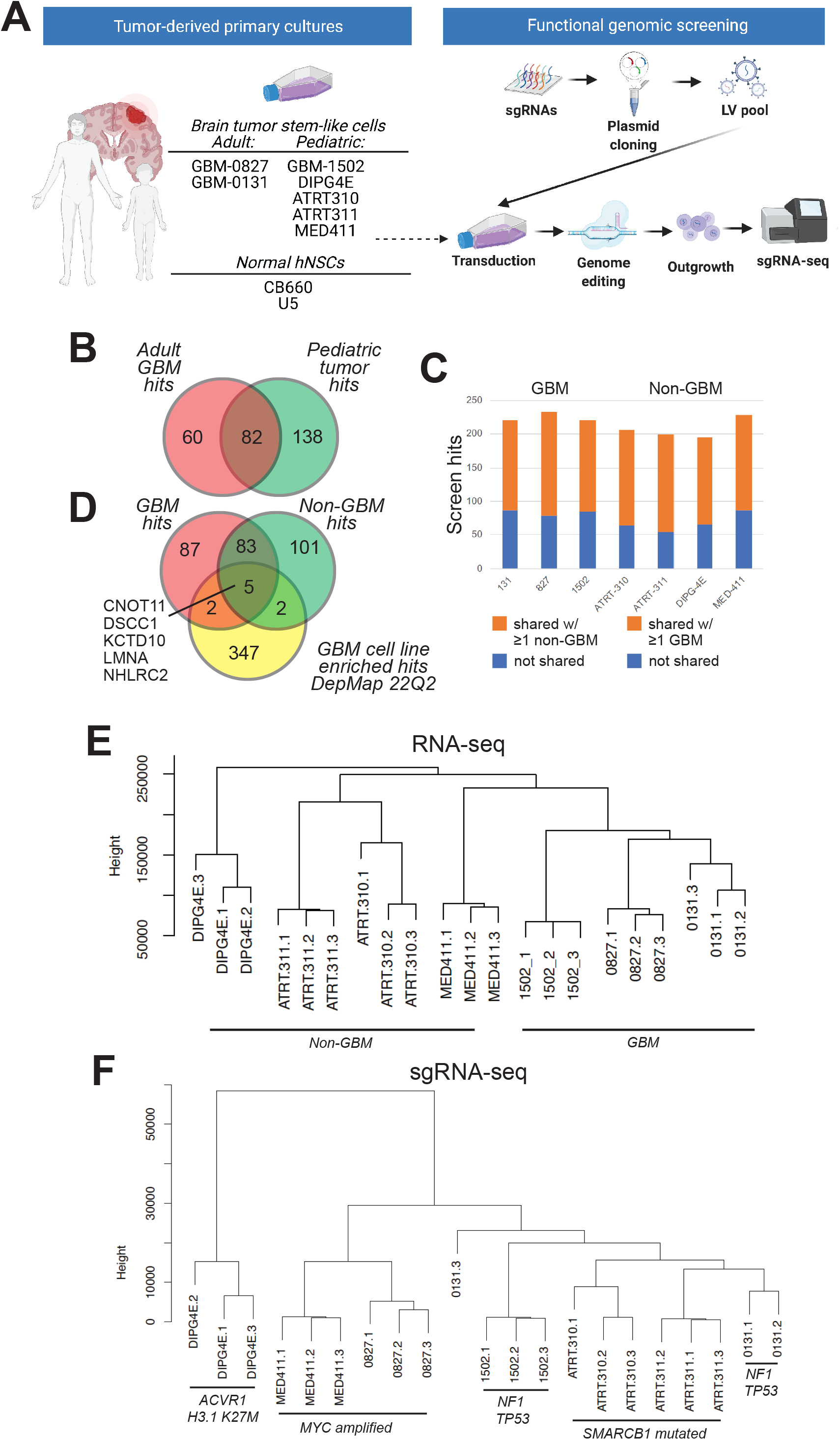
Functional genomic retest lethality screens in adult and pediatric brain tumor isolates. **(A)** Overview of functional genomic screens and data integration. **(B)** and **(C)** Comparisons of scoring screen hits between GBM isolates and non-GBM pediatric brain tumor isolates. **(D)** Comparison of screen data with GBM-enriched cell line screen hits from depmap.org. **(E)** Dendrogram analysis of gene expression analysis for adult and pediatric brain tumor isolates used in comprehensive retest screens used for this study. **Figure S1** show principal component analysis for these same data. Full RNA-seq data are available in **Table S2**. **(F)** Dendrogram analysis of screen results for adult and pediatric brain tumor isolates used in comprehensive retest screens used for this study. **Figure S1** show principal component analysis for these same data. Full screen results are available in **Table S1**.

Retest rates were ~43% for GSC-0131, ~40% for GSC-0827s, and ~53% for GSC-1502s (≥ 2 sgRNAs; Z score <-1, FDR<.05). The frequencies likely reflect a higher false positive rate from the initial screens due to first generation sgRNA designs, relaxed inclusion criteria for retest screens, and the variability of primary cultures. In total, 280 genes validated in one or more brain tumor isolate (**Table S1**).

Overall, ~50% of GBM hits scored in pediatric brain tumors with 46% of 827 hits, 65% of 0131 hits, and 66% of 1502 hits (**Figure 1B**)(**Table S1**). Moreover, the majority of screen hits scoring for each GBM isolates were shared with non-GBM brain tumor isolates and vice versa (**Figures 1C and 1D**). There were no hits in GBM tumors that did not also score in at least non-GBM tumor isolate (**Table S1**).

We also compared lethal gene hits among serum-grown GBM cell line data available via Broad Institute and Sanger Center data (depmap.org). However, we only observe 9 genes overlapping (~3% of total hits) (**Figure 1D**). This is consistent with profound phenotypic divergence observed between serum-free and serum isolated brain tumor cells previously reported [32, 33].

We next compared gene expression and screen data via dendrogram and principal component analysis (**Figures 1E and 1F**; **Supplementary Figures 1B and 1C**). The RNA-seq data reveal isolates cluster by tumor types as GBM or non-GBM tumors (**Figure 1E and S1B**). However, these same relationships were not found among gene dependencies, where pediatric MED-411 clustered with adult GSC-0827 (both share *MYC* amplifications), pediatric isolates from ATRT grouped more closely with adult GSC-0131 cells, and pediatric DIPG-4E cells were an outlier (**Figures 1F and S1C**).

Taken together, these results suggest that, despite dramatic differences in driver mutations, common vulnerabilities likely exist between the adult and pediatric brain tumor isolates, a notion supported by a recently published lethality screens in pediatric cancer cell lines [22].

### Intersection of machine learning models and screening hits reveals candidates for new interactions

We next sought to identify molecular and phenotypic features that predict candidate genetic dependencies. The limited numbers of brain tumor isolates screened were not sufficient to, on their own, create functional genomic networks or associate particular genetic dependencies with genetic features in the patient tumors. Therefore, we endeavored to integrate our lethality screen results with more comprehensive functional genomic lethality screen data sets for >1000 human cancer cell lines now available through the Broad and Sanger Institutes [46, 47]. This included the use of cell line gene effect scores or CERES scores [46]. CERES scores are calculated for every gene in the library and they account for multiple confounding effects that bias the direct measurements of individual sgRNA enrichment and depletion. Negative CERES scores denote net gene depletion and positive scores denote enrichment across sgRNAs.

The Pritchard group has recently incorporated CERES scores with addition cell line features, including (mutation, RNA-seq, CNV, and lineage from the Cancer Cell Line Encyclopedia), to create multivariate machine learning models for gene requirement [43] (**Figure 2A**). This results in thousands of machine-learning models that incorporate the effects of millions of CRISPR knockout phenotypes in addition to mutations, copy number, lineage, and RNA-seq to predict cell type specific phenotypes [43]. For this analysis each screen hit or “target gene” is associated with 10 model features nodes that represent genes that co-vary with requirement for the target gene. The selection of 10 feature models (i.e., multivariate) outperformed single feature models in predicting target gene requirement [43].

**Figure 2:**
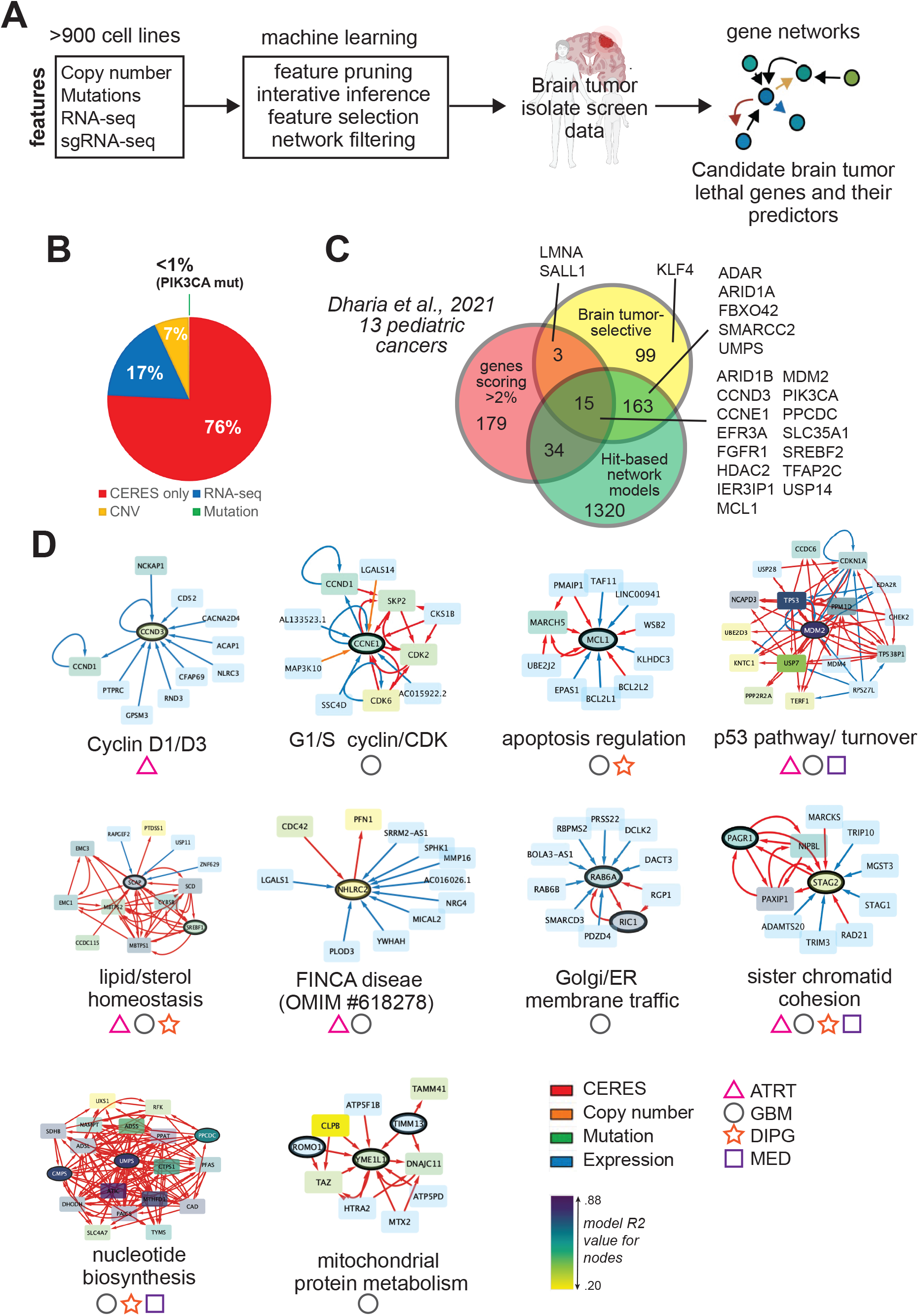
Integration of brain tumor screen data with functional genomic datasets from cancer cell lines. **(A)** Overview of machine learning models used to generate brain tumor screen hit networks and predictive features. **(B)** Proportions of predicted interactions feature prediction ad screen hits. **(C)** Overlap of common essential genes defined by cell line data or **(F)** pediatric cancer dependencies from Dharia et al., 2021 (scoring in >2% of isolates) with all nodes appearing in network models and 280 validating genes scoring in 1 or more brain tumor isolate. **(D)** Examples of first neighbor networks from **Table S3**. Networks are centered on primary screen targets: central node (oval, black boarders). Other screen hits are also shown as oval nodes with black boarders.

For our data sets, we used 280 screen hits to populate cell line network models and identify candidate predictive features associated with genetic dependencies (**Table S3**). 76% or 1348 features were based on CERES score interactions, while 17% or 309 were based on gene expression, 7% or 121 CNV and <1% or 1 mutation (**Figure 2B**). This is in-line with assessment of all cancer cell line network models, where 73% of predictive features are based on CERES scores (i.e., co-requirement for the two genes) 22% on RNA-seq, 5% on CNV, and <1% on mutation [43]. Thus, most network interactions exist as “co-dependencies”, where requirement for the target gene tends to co-occur with requirement for feature genes (i.e., CERES score-based edges).

Comparisons of network features from all cancer cell line data, common essential genes, and our validating screen hits revealed there were 190 screen hits that are shared among the total network nodes and 71 genes which are brain tumor selective (i.e., not essential or represented in networks) (**Figure 2C**). A portion of our screens hits and network work features contain common essential genes (~31% of validating screen hits). Inclusion of these genes (e.g., TATA binding protein (TBP)) reflects how these scored in brain tumor isolates relative to NSCs both in the original screen and validation screens.

Gene set enrichment analysis for all brain tumor network nodes revealed enrichment for genes associated with cell cycle, p53 pathway, chromatin modification, DNA damage response, and mTOR pathway (**Figure S2**). They were also enriched in genes associated with other cancers such as small cell lung and prostate cancer.

Comparisons to recently published functional genetic dependencies in pediatric cancer cell lines [22] showed 18 candidate dependencies directly in common and 49 overall that appear in hit-based feature models and have feature predictions (**Figure 2C**). These included, for example, MCL1 gene dependency, which were associated with BCL2L1 expression and which our models also predict, as well as EFR3A, FGFR2, MDM2, PIK3CA, etc. (**Figures 2C and 2D**). The portion of pediatric dependencies not shared likely include dependencies from non-CNS cancers and false negatives from our library gene selection process.

Among the results, we find previously identified functional genetic interactions and predictive associations in our isolates. These include MDM2 requirement in p53/TP53 wildtype cancers [48]. As shown in Figure 2B, MDM2 is differentially required where p53 protein is functional among the tumor isolates; these include ATRT310, ATRT311, GSC-0827, and MED411 isolates, which are not mutated for TP53 *and* sgTP53 causes growth enhancement. The predictive MDM2 network includes TP53 other known p53 transcriptional targets including CDKN1A [49] and RPS27L [50] (**Figure 2D**).

Other examples include ARID1A and ARID1B synthetic lethality (by CERES score)[27], same gene interactions for CCND3 and CCNE1 (by RNA-seq) [30], paralogous functional redundancy between RAB6A and RAB6B [30] and ZFY and ZFX (by RNA-seq)[51], and association of interferon gene expression with ADAR requirement (by RNA-seq)[52].

The sole mutational predictor among our screens hits was PIK3CA, where hotspot mutations predict requirement for PIK3CA. PIK3CA did not score in the original GBM lethal screens but was included in the retest pool as a biological control. It scored significantly only in DIPG4E cells (**Table S1**). Examination of RNA-seq reads across PIK3CA locus revealed that DIPG4E cells indeed harbor a G to A mutation in a preponderance of RNA-seq reads at codon 545 which changes glutamate to lysine, an activating, hotspot mutation found DIPGs and other cancers [53](not shown).

Given these confirmatory results, we explored four gene dependencies identified in network models for our brain tumor isolates and their possible implications for brain and other tumors.

### EFR3B expression predicts EFR3A requirement

EFR3A and EFR3B are paralogs (ensembl.org) that function redundantly to localize an adapter protein required to localize PI4KIIIα to the plasma membrane and regulate PtdIns(4)P synthesis [54, 55] (**Figure 3A**). EFR3A scored as a screen hit in GSC-0131 cells, but not other isolates. As predicted by network model and supported by cell line screening data, GSC-0131 cells show lower expression of EFR3B expression and a requirement for EFR3A (**Figures 3B and 3C**). Ectopic expression of EFR3B suppressed requirement for EFR3A alone in GSC-0131 cells or EFR3A and EFR3B together in GSC-0827 cells (**Figures 3B and 3C**). Analysis of EFR3B expression in wide variety adult and pediatric tumors and control brain tissues revealed that its expression is significantly reduced (compared to control GTEX brain samples) in adult GBM (IDH-wt) (p<.001; n= 348) and the following pediatric tumors: ATRT (p<.01; n=30), meningioma (p<.001; n= 29), neurofibroma (p<.001; n= 21), and Schwannoma (p<.001; n= 19) tumors (**Figures 3F and G**) (**Table S4**). These results suggest that loss of EFR3B expression predicts sensitivity to its paralog EFR3A. The ancestral EFR3 gene is predicted to have duplicated ~796 MYA in bilateral animals (Ensemble GeneTree ENSGT00390000002143). However, yeast still harbor a single EFR3 gene which, similar to EFR3A/B function in mammals, is essential for recruiting PtdIns 4-kinase to plasma membrane [56].

**Figure 3:**
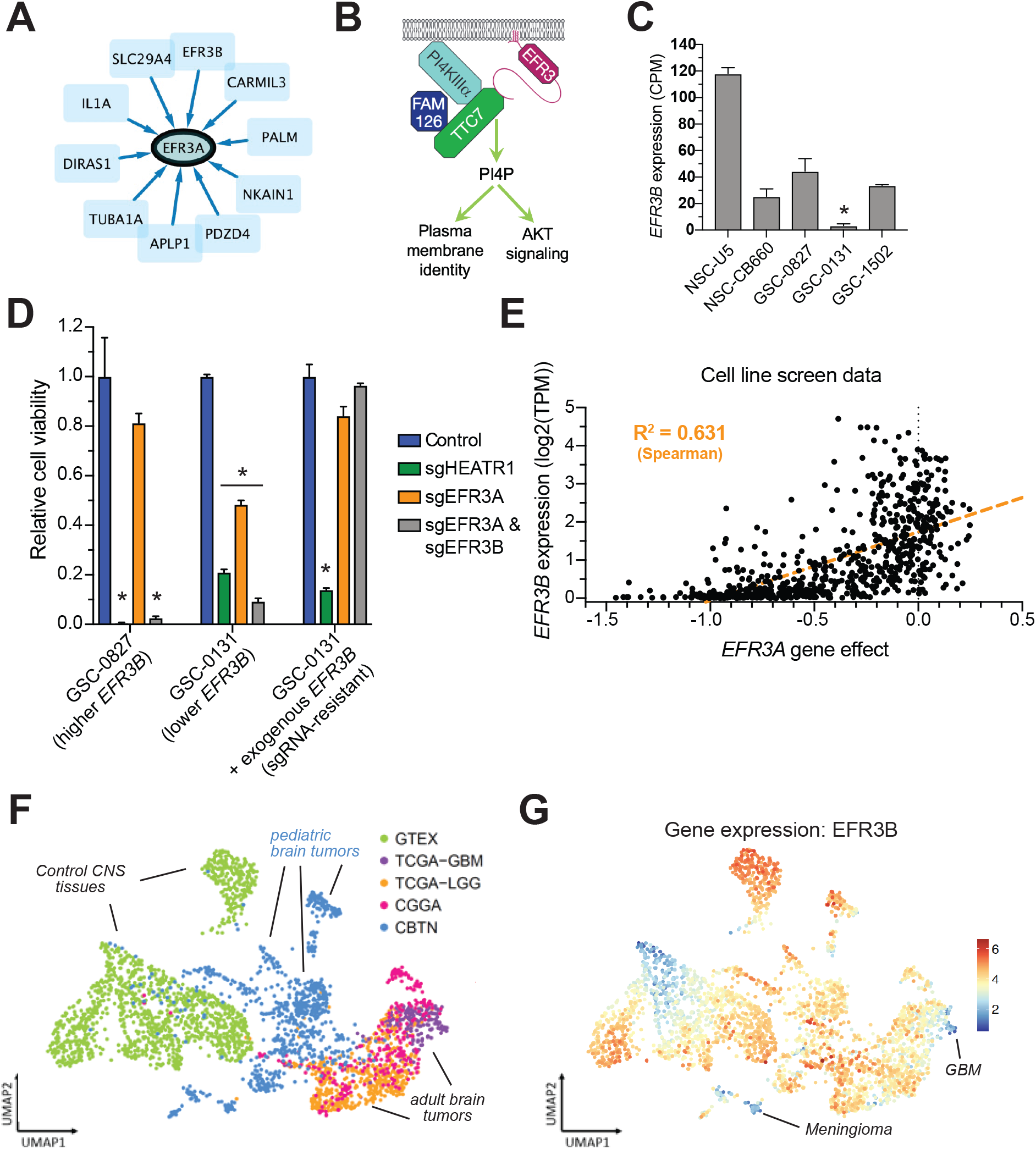
EFR3B expression predicts EFR3A requirement. **(A)** First neighbors network for EFR3A predicting that EFR3B expressing predicts requirement for EFR3A (key is same as **Figure 2**). **(B)** Cartoon of EFR3A/B adaptor function. **(C)** Expression of EFR3B from RNA-seq data (n=3) (NSC = neural stem cell). **(D)** Relative cell viability (normalized to targeting control sgCD8A) for cells nucleofected with CRISPR RNPs targeting EFR3A, EFR3B, or both. GSC-0131 are compared to GSC-0131 that were transduced with a lentiviral construct expressing EFR3B. HEATR1 is an essential control gene. sgNTC = non-targeting control sgRNA. Measured at 9 days post nucleofection. **(E)** Cell line functional genomic screening data showing EFR3A dependency score vs. EFR3B expression. Each dot represents a cell line. Dotted blue line shows linear regression fit. **(F) & (G)** UMAP projections of “bulk” gene expression data from brain tissue and tumor samples. **(F)** UMAP projection of gene expression showing sample source. Tumors and tissue samples colored by study. CBTN= Children’s brain tumor network; CGGA= Chinese Glioma Genome Atlas (adult); GTEX= The Genotype-Tissue Expression Project; HGG= High grade glioma (adult); LGG= Low grade glioma (adult); TCGA = The Cancer Genome Atlas. Tumor type break down can be found in **Supplementary Figure S3**. **(G)** UMAP projection of gene expression for EFR3B for **(F).** Scale bar is Log2(TPM+1). A key for all tumors in this plot is available in **Figure S3** and **Table S4**. *indicates p<.01, student’s t-test.

### FGF2 predicts requirement for FGFR1

Similar to PIK3CA, the receptor tyrosine kinase gene FGFR1 was added as a biological control in validation screens given its importance in regulating developmental potential of neural stem/progenitor cells [32, 57]. Among our isolates, pediatric ATRT and DIPG cells showed specific dependency on FGFR1 for outgrowth (**Figure 4A**). The FGFR1 feature network reveals multiple co-dependencies by CERES score of genes required for FGFR1 signaling (**Figure 4B**). These include FRS2 expression, which encodes a key FGFR1 target that regulates neural stem/progenitor cells self-renewal [58], and genes required for synthesis of heparan sulfate, a mandatory cofactor in paracrine FGF signaling [59], including B3GAT3, EXT1, EXT2, EXTL3, GLCE, HS2ST1 and SLC35B2 (Reactome database).

**Figure 4:**
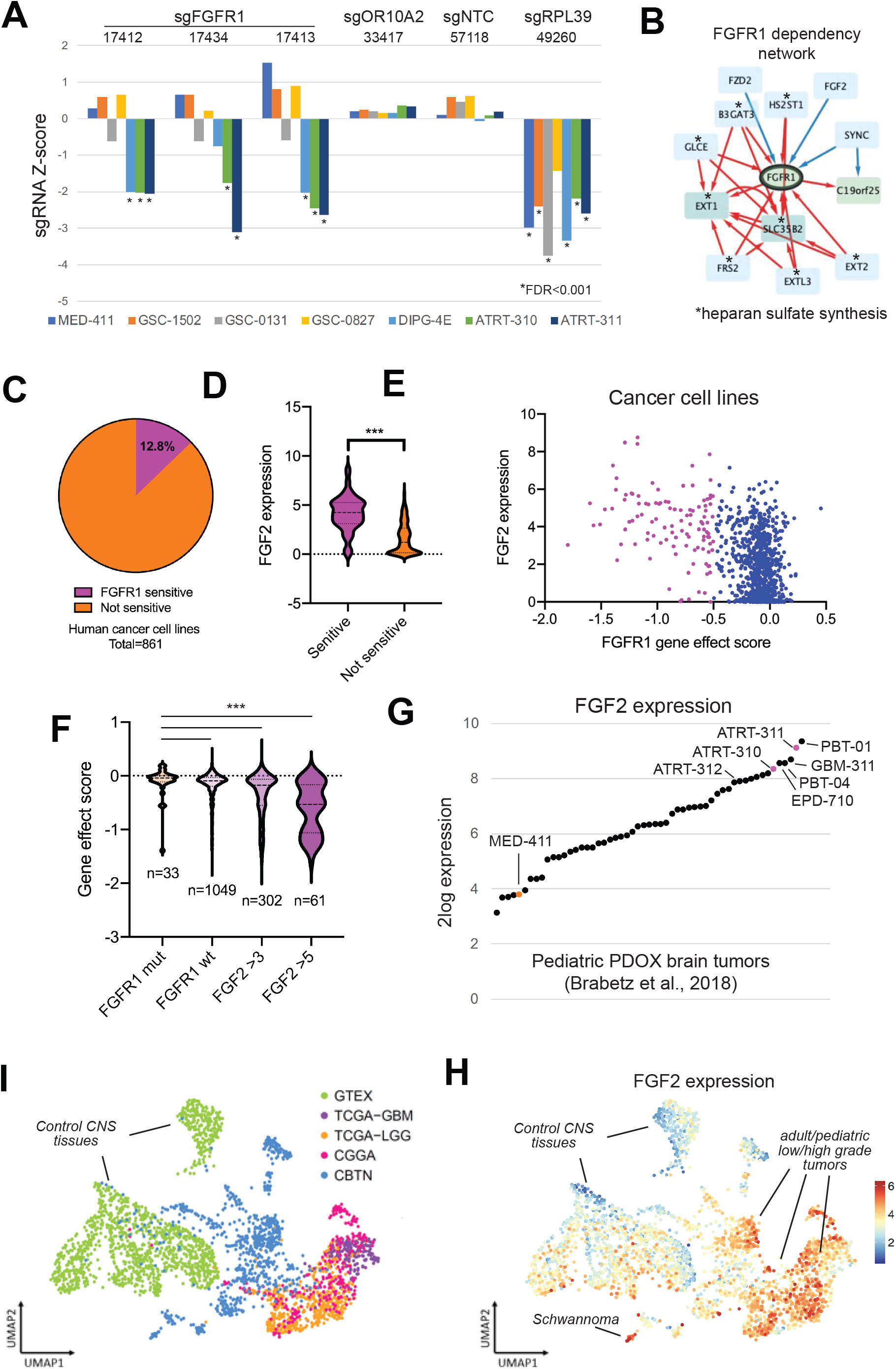
FGF2 expression predicts FGFR1 requirement. **(A)** Retest results of 3 FGFR1 sgRNAs that score differentially in ATRT and DIPG isolates. Controls include an sgRNA targeting olfactory receptor gene OR10A2 (nonessential), a non-targeting control (NTC), and one targeting RPL39, a common essential gene. **(B)** First neighbors network for FGFR1, where FGF2 expression predicts requirement for FGFR1 (key is same as **Figure 2**). **(C-E)** Data derived from Broad and Sanger cell line screens. **(C)** Sensitivity to FGFR1 loss defined by CERES score (<-0.5 = sensitivity; >-0.2 = insensitivity). **(D)** FGF2 expression in units of log2 transcript count per million (TPM) for FGFR1 sensitive and insensitive cell lines. Mann Whitney U test was used to test significance. *indicates p<.001. **(E)** FGF2 expression in units of log2 TPM versus FGFR1 CERES score for all Sanger and Broad cell lines. Lines with CERES score <-0.5 are highlighted. **(F)** Comparisons of CERES scores for cell lines with FGFR1 mutations, FGFR1 wt, FGF2 expression >3 log2TPM, and FGF2 expression >5 log2TPM. Mann Whitney U test was used to test significance. *indicates p<.001. **(G)** FGF2 expression from patient-derived orthotopic xenograft (PDOX) models of pediatric brain tumors (Brabetz et al., 2018). **(H)** UMAP projections of gene expression data for FGF2 from brain tissue and tumor samples. Key for samples is same as **Figures 3F and S3**. Scale bar is Log2(TPM+1). **(I)** Gene expression analysis of FGF2 adult and pediatric brain tumors and normal brain tissues from CBTN, CGGA, GTEX, and TCGA databases. Tumor lists and statistical test data are available in **Table S4**. *indicates p<.001. Green indicates significant reduction in expression.

The expression of FGF2, one of FGFR1’s key ligands [60], also scored as being predictive of FGFR1 requirement (**Figure 4B**). This was true for cancer cell lines derived from diverse tumor types: bone, brain, breast, liver, lung, skin, and soft tissue (**Figure 4C-E**). Moreover, FGFR1 mutational status was not associated with sensitivity, when compared to FGFR1 wt or FGF2 high expressors (**Figure 4F**). 33 out of 1082 cell lines are shown in **Figures 4C-E** are FGFR1 mutant; only 3 of 33 score as effect score <-0.5. Examination of FGF2 expression among our FGFR1-sensitive and insensitive brain tumor isolates supported this notion. ATRT-310, ATRT-311, and DIPG4E, show higher expression of FGF2 than the insensitive cells *in vitro* (**Figure S4A**) and *in vivo* (**Figure 4G**). Analysis of tumor expression of FGF2 revealed most adult low and high grade gliomas and 12 of 18 pediatric tumor types examined (e.g., DIPG, ependymoma, pilocytic astrocytoma, etc.) have significantly higher expression than normal tissue controls (**Figures 4H** and 4I)(**Table S4**). By contrast, pediatric medulloblastoma and meningioma showed significantly less FGF2 expression (**Figure 4I**). Thus, FGF2 expression is a candidate biomarker for sensitivity to FGFR1 network inhibition.

### SMARCB1 mutant ATRT tumor isolates show requirement for SMARCC2

SMARCC2 was included in our retest library due to its effect in GSC-0827 screens, where a single sgRNA scored using relaxed inclusion criteria. SMARCC2 codes for a non-essential core subunit of the SWI/SNF chromatin remodeling complex core module and is structurally redundant with SMARCC1 [61]. SMARCC2 is part of a SWI/SNF complex dependency network and is associated with requirement for other SWI/SNF complex members, including ARID1A, ARID1B, DPF2, SS18, SMARCB1, SMARCC1, and SMARCE1 (**Figure 5A**) [61]. However, Figure 5B shows that SMARCC2 is specifically lethal among ATRT isolates, scoring similar to common essential gene hits. SMARCC2 KO did not significantly growth affect the growth of U5 or CB660 NSCs (**Table S1**).

**Figure 5:**
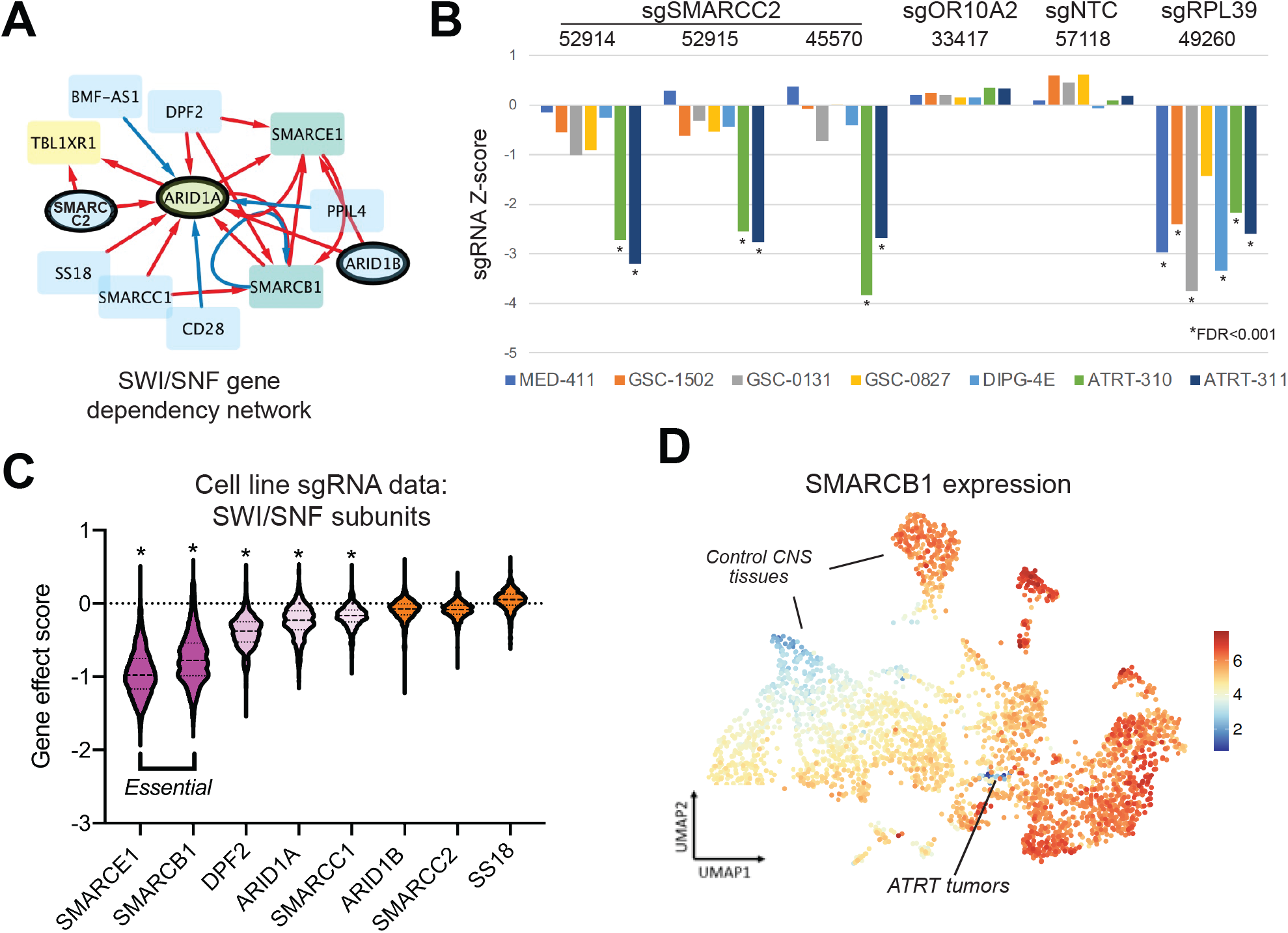
Examination of SMARCC2 requirement. **(A)** First neighbors network for ARID1A revealed associations with ARID1B, SMARCC2, and also SMARCB1. **(B)** Retest results showing SMARCB1 deleted/mutated ATRT isolates specifically require SMARCC2. Controls include an sgRNA targeting olfactory receptor gene OR10A2 (non-essential), a non-targeting control (NTC), and one targeting RPL39, a common essential gene. **(C)** Comparisons of gene effect scores among SWI/SNF subunits found in (A). *indicates p<.001, Mann Whitney U test, where gene effect scores for shown gene are compared to SMARCC2 scores. **(D)** A UMAP projection of SMARCB1 expression in adult and pediatric brain tumors. Key is available in **Figure S3** and **Table S4**. Scale bar is Log2(TPM+1). **(E)** Gene expression analysis of SMARCB1 in adult and pediatric brain tumors and normal brain tissues from CBTN, CGGA, GTEX, and TCGA databases. Tumor lists and statistical test data are available in **Table S4**. *indicates p<.001.

~95% of malignant rhabdoid tumors, which include ATRTs, have biallelic inactivation of SMARCB1 [62, 63], a key component of “canonical” BAF and PBAF complexes [61]. While SMARCB1 is thought of as a is a tumor suppressor for rhabdoid and certain other cancer types, it is, none the less, essential in proliferating somatic cell lines as shown in the cell line screen data (as is another key component SMARCE1) (**Figure 5C**). By contrast, SMARCC2 is non-essential in somatic cell lines (Figure 5C, possibly owing to its structural redundancy with SMARCC1 [61].

Further, SMARCC2 and SMARCB1 dependency co-occurred in our SWI/SNF network model (**Figure 5A**). However, SMARCB1 mutation or expression was not associated with SMARCC2 requirement. This is likely due to underrepresentation of rhabdoid tumors among cancer cell line data (e.g., n=5 ATRT cell lines) and overrepresentation of the SMARCB1 sensitive lines.

We also examined SMARCB1 gene expression across multiple types of adult and pediatric brain tumors (**Figures 5D and 5E**)(**Table S4**). This analysis revealed that loss of SMARCB1 expression is highly selective for ATRT tumors (n=30), consistent with biallelic inactivation (**Figure 5E**). However, meningiomas also showed significant reduction (p<.001; n=29) (**Table S4**) and ganglioglioma and schwannomas showed a tendency (p<.16).

This data suggests that SMARCC2 is required in the context of SMARCB1-deficient ATRT tumors and that gene expression is likely a suitable biomarker of loss of SMARCB1 function.

### ADAR requirement in adult and pediatric brain tumor isolates

ADAR was originally a screen hit in adult GSC-0131 cells [41] that validated in both GSC-0131 and DIPG4E cells (**Figure 2B**). ADAR, also known as ADAR1, is a dsRNA editing enzyme that post-transcriptionally converts adenosine-to-inosine in both coding gene mRNAs and repetitive genomic element RNAs [64]. It was recently shown that tumor cells displaying interferon-stimulated gene (ISG) expression signatures require ADAR activity to prevent accumulation of cytotoxic dsRNA species [52, 65]. The network associated with ADAR gene dependency fits well with this concept (Figure 6A). It includes many ISG genes (e.g., MX1) as predictive RNA-seq based features that are negatively correlated and more highly expressed in cells with ADAR requirement. Consistent with this notion, sensitive brain tumor isolates and in general a portion of adult and pediatric brain tumors have increased ISG gene signature (Figures 6B and S6A).

**Figure 6:**
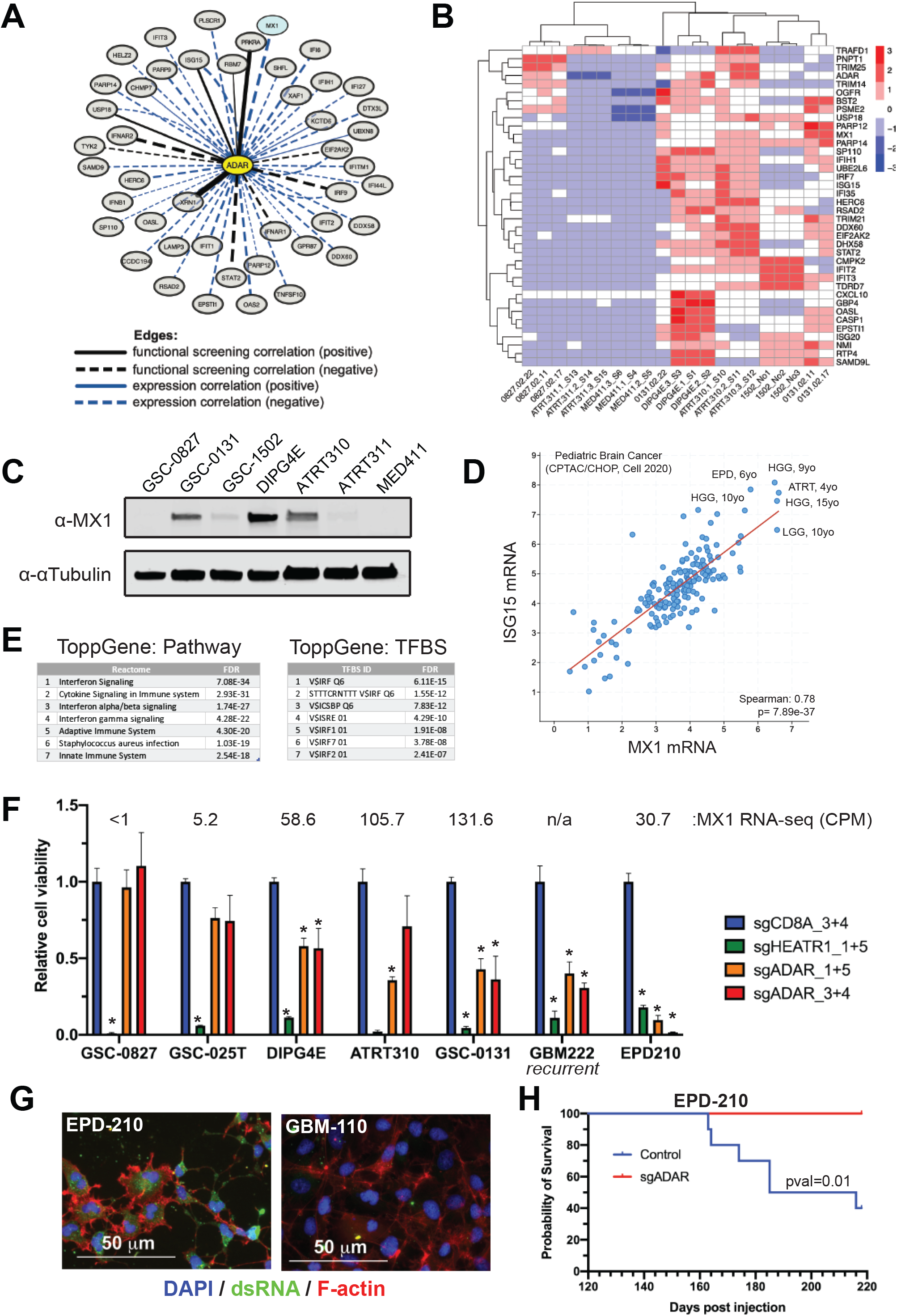
Validation of ADAR requirement in adult and pediatric brain tumor isolates. **(A)** Mini-network of top gene co-dependency and gene expression correlations for the GSC-0131-specific screen hit ADAR. Blue expression edges indicate that expression of the gene to which ADAR is connected correlates with ADAR dependency score. **(B)** Heatmap of gene expression analysis of screened brain tumor isolates for a 38-gene IFN-stimulation signature (from [52]). IFN signature among pediatric brain tumor isolates is available in **Figure S5**. **(C)** Western blot showing MX1 protein expression in brain tumor isolates screened. Additional examples are shown in Figure S6. **(D)** Comparison of MX1 and ISG15 expression in pediatric brain cancers (expression of both are associated with ADAR requirement in **(A)**). Additional MX1 expression in brain tumors can be found in **Figure S5**. **(E)** Pathway and transcription binding factor analysis of top 200 genes positively correlated with MX1 expression among pediatric cancers shown in (D) from [93]. Analysis was performed using ToppGene [94]. **(F)** Relative cell viability (normalized to targeting control sgCD8A) for lines nucleofected with CRISPR RNPs targeting ADAR. HEATR1 is an essential control gene. sgNTC = non-targeting control sgRNA. Measured at 10 days post nucleofection (n=3; *pval.<.01 Student’s t-test). **(G)** Immunofluorescence for dsRNA of pediatric isolates EPD-210 (MX1 expressed; ADAR sensitive) and GBM-110 (MX1 negative). **(H)** Survival of mice with EPD-210 tumors with control sgRNA or sgADAR. Log-rank (Mantel-Cox) test was used to evaluate significance (n=8 for each arm).

To confirm these results, we assayed protein expression of several ISG genes, including MX1, ADAR, OAS1 and IRF9 (**Figures 6C, S5C and S5D**). Of these, MX1 produced a robust signal which appears to correlate with ADAR sensitivity in screen lines and also other isolates (**Figures 6C, S5D and S5E**). These included a pediatric ependymoma and two matched, recurrent GBM isolates.

Since ADAR gene dependency has not been established in pediatric brain tumors, we next examined genes that are enriched with MX1 expression in pediatric brain cancers (**Figures 6D and 6E**). Not surprisingly, ISG genes were among the top genes correlated with MX1 expression, which included ISG15. Top pediatric cancer expressors ISG15 and MX1 included ATRT, EPD, and low and high grade gliomas (**Figure 6D**). Moreover, we could find statistically significant increase in MX1 expression in most adult and pediatric brain tumor types surveyed, with the exception of pediatric embryonal tumors, medulloblastomas, and DNETs (**Table S4**).

Further retests of ADAR KO in these isolates displayed a range of *in vitro* sensitivities where were in line with MX1 expression, where recurrent GBM-222 and EPD-210 had the greatest loss of viability (**Figure 6F**). EPD-210 cells harbor high levels of dsRNA relative to GBM-110 non-MX1 expressing, a hallmark of ADAR sensitive [52] (**Figure 6G**). Further, inhibiting ADAR in the context of orthotopic EPD-210 brain tumors had a significant survival benefit (**Figure 6H**). Taken together, these data confirm that ADAR gene dependency in variety of adult and pediatric brain tumors and is predicted by MX1 protein expression.

## Discussion

There is currently a critical need for new therapeutic strategies for adult and pediatric brain tumors. A primary goal of this study was to determine whether functional genetic relationships from cancer cell line data, mostly derived from non-CNS cancers, could be leveraged to reveal and characterize gene dependencies in adult and pediatric brain tumors. Our results reveal that this integrated approach can be powerful tool for discovery of candidate cancer liabilities and their molecular predictors in patient tumor isolates.

In-line with other studies, we found that most gene dependencies in brain tumor isolates are predicted by other gene dependencies, rather than gene expression or genetic alterations. The lone mutational predictor was a PIK3CA hotspot mutation predicting PIC3CA requirement in DIPG cells (**Table S2**). Instead, our results show that brain tumor networks models can illuminate connections between candidate genetic vulnerabilities and candidate biomarkers in brain tumors, capturing previously described and novel ones. In total, we provide machine learning networks with candidate predictors for 280 brain tumor screen hits (**Figure 2; Table S2**).

EFR3B expression predicts sensitivity to its paralog EFR3A (**Figure 3**). These genes function redundantly in other systems to localize PI4KIIIα to the plasma membrane and regulate PtdIns(4)P synthesis [54, 55]. EFR3B expression is enriched in brain, muscle, and endocrine tissues and is associated with neuronal signaling, while EFR3A is expressed in most tissues (www.proteinatlas.org). EFRB3 expression is lost in a subset of brain tumor types including GBM and multiple pediatric tumors (e.g., ATRT, meningioma, neurofibroma and Schwannoma). The results suggest that EFR3A or PtdIns(4)P synthesis is worthy of consideration as a selective vulnerability in EFR3B low tumors.

FGF2 expression predicts sensitivity to FGFR1 inhibition. FGFR inhibitors are currently in clinical trials for a variety of tumors, including GBM (e.g.,[66]). However, these trials tend to focus exclusively on FGFR-altered tumors with mixed results. Both ATRT and DIPG tumor isolates were sensitive to FGFR1 loss but are not mutated for FGFR1 (**Figure 4**). Instead, the FGFR1 dependency network suggested that FGF2 expression is associated with FGFR1 requirement. Consistent with this, ~13% of cancer cell lines that show sensitivity to FGFR1 loss have significantly higher FGF2 expression (**Figures 4C-E**). FGFR1 mutant cells, on the other hand, were significantly less likely to be FGFR1 sensitive, compared to wt or FGF2 high cells. Thus, the results imply that FGF2 expression maybe a valid indicator of FGFR1 sensitivity in FGFR wt cancers.

We observed a novel requirement for SMARCC2, a core SWI/SNF subunit, in ATRT isolates (**Figure 5**). Mutant rhabdoid tumors, including ATRTs, have biallelic inactivation mutations in SMARCB1 [62, 63], which codes for a key “canonical” SWI/SNF complex subunit [61]. In most other cancer cell types, however, SMARCB1 is essential; by contrast, SMARCC2 is not (**Figure 5C**). Previous studies have shown that SMARCB1 mutant rhabdomyosarcoma and synovial sarcoma cells are sensitivity to loss of “non-canonical” BAF components, including BRD9 and GLTSCR1, to maintain cancer-specific chromatin structure and gene expression [67, 68]. Because ATRT tumors are predicted to be deficient in canonical SWI/SNF function, novel SMARCC2 requirement may arise due to its participation in noncanonical SWI/SNF complex [61]. For example, ncBAF includes a BAF core consisting of SMARCC1/2, SMARCD1/2/3) and an ATPase module (SMARCA2/4) but not other commonly studied SWI/SNF complex proteins (e.g., ARID1A/B, DPF2, SMARCB1, SMARCE1, etc.) [61]. Thus, ATRTs and also possibly meningiomas (**Figure 5E**) may show differentiation requirement for ncBAF, which may be an actionable target (e.g., via BRD9 inhibition [69] or targeted degradation [70]). However, future studies will need to demonstrate gene dependencies for other ncBAF components.

We also found that ADAR requirement in brain tumors is predicted by interferon gene expression and MX1 protein expression (**Figure 6**). Previous studies have suggested that ADAR influences glioma cell proliferation either through regulation of the m^6^A-methyltransferase activity [71] or translation control [72]. However, our results and network model predictions are in-line with reports in other cancers, whereby interferon signaling is triggered by the aberrant accumulation of dsRNA [52] (**Figure 6G**). MX1 expression is significantly increased in most adult and pediatric brain tumor types surveyed compared to normal brain tissue. One such tumor, MX1 high pediatric ependymoma, was shown to be sensitive to ADAR loss *in vivo* (**Figure 6H**). Additionally, we find that recurrent GBM isolates can display high MX1 expression, consistent with proneural-to-mesenchymal transition after SOC [73], and are sensitive to ADAR loss. Thus, ADAR may be a selective target for a broad range of brain tumors.

## Limitations

Basing networks on cancer cell line data from current functional genomic libraires has a few key limitations. First, the cellular heterogeneity found in tumors (e.g., [74, 75]) is missed. GBM tumors, for example, contain cell populations with diverse neuro-developmental states that co-exist in tumors (e.g., [74, 75]), which may contribute to their recurrence. One relevant example is the mesenchymal cell state, which is associated with interferon gene expression and ADAR requirement (Figure 6). Targeting ADAR in tumors would likely affect only the mesenchymal subpopulations. Similarly, many brain tumor-specific gene dependencies are not found among cell line data (Figure 2E). Such limitations may be addressed by using new computational strategies to reduce CRISPR library complexity while retaining predictive power [43] for screens in primary tumor systems *in vivo*.

Other limitations include the absence of paralogous gene interactions, which require two or more genes to be targeted simultaneously [76], and the absence of noncoding RNAs and other genetic elements not present in screening libraries.

## Supporting information

Supplemental Inventory and Figures

## Acknowledgements

We thank members of the Paddison and Pritchard labs and Adam Geballe for helpful discussions, and Pam Lindberg and An Tyrrell for administrative support. This work was supported by the following grants: Interdisciplinary Training in Cancer Fellowship NCI T32CA080416 (P.H.); NCI/NIH (R01CA190957) (P.P.) (R01CA114567) (J.O.); and NINDS/NIH (R01NS119650) (P.P.). This research was funded in part through the NIH/NCI Cancer Center Support Grant P30 CA015704.

## Author Contributions

Project conception and design was carried out by P.H., M.B., J.H., J.O., J.P., and P.P.; experiments and data analysis were performed by P.H., M.B., E.G., P.D., K.M., and D.K.; critical reagents were generated by M.K., L.S., and J.O.; bioinformatical data analysis and statistics were performed by P.H., S.A., and Y.R. with input from J.P., E.H., and P.P.; P.P. and P.H. wrote the manuscript with input from other authors. Funding acquisition: E.C., J.O., J.P., and P.P.

## Methods

*Key Reagents and Resources are available in Table S6*.

### Cell Culture

Isolates were cultured in NeuroCult NS-A basal medium (StemCell Technologies) supplemented with B27 (Thermo Fisher Scientific), N2 (homemade 2x stock in Advanced DMEM/F-12 (Thermo Fisher Scientific)), EGF and FGF-2 (20 ng/ml) (PeproTech), glutamax (Thermo Fisher Scientific), and antibiotic-antimycotic (Thermo Fisher Scientific). Cells were cultured on laminin (Trevigen or in-house-purified)-coated polystyrene plates and passaged as previously described [77], using Accutase (EMD Millipore) to detach cells.

### Primary GBM cultures

Primary, patient-derived GBM cultures were generated in-house from tissue samples obtained during surgical resection of patients diagnosed with GBM. As previously described [78], tumors were subjected to enzymatic digest, mechanically dissociated and cultured as neurospheres. GBM neurospheres were expanded as intracranial xenografts (PDX) in athymic nu/nu mice and processed as previously described by [79]. Cultures from PDX samples were processed similarly to the patient derived tumors. Freshly isolated human GBM samples were obtained under an IRB approved protocol (approval SHIRB # 2015.059-1).

### CRISPR-Cas9 library

For generating our comprehensive retest library, sgRNAs for chosen genes and controls were mined from the human GeCKO v2 library [80] as well as our whole genome screen results, and an oligo pool representing these sgRNAs and PCR adapters was obtained (Twist Bioscience). The oligo pool was PCR amplified using Herculase II Fusion DNA Polymerase (Agilent) and cloned into lentiCRISPRv2 puro vector (Addgene) using Gibson Assembly Master Mix (New England Biolabs). The assembled pool was then transformed into Stellar Competent Cells (Clontech) and plated onto LB agar plates (liquid culture was avoided in order to minimize competition between clones containing different sgRNAs). The resulting colonies were scraped from the plates and the finished lentiCRISPRv2 plasmid comprehensive retest library was extracted using a NucleoBond Xtra Midi Endotoxin-Free Kit (Macherey–Nagel). This lentiviral library was then used to generate virus and infect cells for outgrowth screening.

### Lentiviral Production

For virus production, lentiCRISPR v2 plasmids [81] were transfected using polyethylenimine (Polysciences) into 293T cells along with psPAX and pMD2.G packaging plasmids (Addgene) to produce lentivirus. For the whole-genome CRISPR-Cas9 libraries, 25×150mm plates of 293T cells were seeded at ~15 million cells per plate. Fresh media was added 24 hours later and viral supernatant harvested 24 and 48 hours after that. For screening, virus was concentrated 1000x following ultracentrifugation at 6800*xg* for 20 hours. For validation, lentivirus was used unconcentrated at an MOI<1.

### CRISPR-Cas9 Screening

For screening, cells were transduced to achieve ~750X representation of the library (at ~30% infection efficiency to ensure a high proportion of single integrants). 2 days after transduction, media was replaced with media containing 2 μg/mL puromycin. After 3 days of selection, portions of cells representing 500-750X coverage of the library were collected as the “Day_0” samples. The remaining cells were cultured and consistently maintained at 500-750X representation for 21-23 days, after which time the “Day_final” samples were collected. Screening was carried out in triplicate. To read out screen results, genomic DNA was extracted using the QIAamp DNA Blood Mini Kit (QIAGEN), and a two-step PCR procedure was used to first amplify the genomically integrated sgRNA sequences and then to incorporate Illumina deep sequencing adapters and barcodes onto the sgRNA amplicons. For the first round of PCR, a sufficient number of PCR reactions were carried out to use all gDNA from the 500-750X coverage sample of cells at 2μg genomic DNA per PCR reaction, using MagniTaq Multiplex PCR Master Mix (Affymetrix) and 12 cycles. For the second round of PCR, 5μL of the first round product was used as a template in combination with primers that would add the deep sequencing adapters and barcodes, using Herculase II Fusion DNA Polymerase (Agilent) and 16 cycles. Amplicons from the second round PCR were then column purified using the PureLink Quick PCR Purification Kit (Invitrogen). Purified PCR products were sequenced using HiSeq 2500 (Illumina). Bowtie [82] was used to align the sequenced reads to the sgRNA library, allowing for 1 mismatch. The R/Bioconductor package edgeR [83] was used to assess changes across groups.

### RNA-seq analysis

Cells were lysed with Trizol (Thermo Fisher). Total RNA was isolated (Direct-zol RNA kit, Zymo Research) and quality validated on the Agilent 2200 TapeStation. Illumina sequencing libraries were generated with the KAPA Biosystems Stranded RNA-Seq Kit[84] and sequenced using HiSeq 2000 (Illumina) with 100bp paired-end reads. RNA-seq reads were aligned to the UCSC hg19 assembly using STAR2 (v 2.6.1)[85] and counted for gene associations against the UCSC genes database with HTSeq [86]. Normalized gene count data was used for subsequent hierarchical clustering (R package ggplot2 [87]) and differential gene expression analysis (R/Bioconductor package edgeR [83]). Heatmaps were made using R package pheatmap [88].

### Model building and network analysis

The retrieval and preprocessing of Cancer Dependency Map datasets 19Q3 and 19Q4 data were performed as in [43]. Features (CERES score, RNA-seq, copy number, mutation, lineages) derived from multiple omics datasets for 18333 genes were used to build predictive models for each of the 280 brain tumor screen hits in the Broad data. The model building process is provided at the Pritchard Lab at PSU GitHub repository [https://github.com/pritchardlabatpsu/cga]. Briefly, a regression model with iterative random forest and Boruta feature selection was used to select each target brain screen hit gene’s top 10 predictive features. The top 10 features were then used to fit a new reduced feature random forest model. To evaluate the model, only qualified gene models(R2>0.1, recall>0.95, defined in [43]) were selected. This left 178 predictable genes and their top 10 features used in subsequent analyses. Default parameters were used in the model building process: random forest regressor:1000 trees; maximum depth:15 per tree; minimum samples required per leaf node: 5; maximum number of features:log2 of the total number of features.

### Network analyses

Networks were derived from the model results in [43]. A fully documented git repository with all source codes and notebooks can be accessed at the Pritchard Lab at PSU GitHub repository [https://github.com/pritchardlabatpsu/cga]. The data used as the input to this study are available in the Zenodo database under the DOI identifier (10.5281/zenodo.5721869). Network communities surrounding hits (N=1 nearest neighbors in the published network) were extracted and visualized using Cytoscape v3.7.256.

### Western Blotting

Cells were harvested, washed with PBS, and lysed with modified RIPA buffer (150mM NaCl, 25mM Tris-HCl (pH 8.0), 1mM EDTA, 1.0% Igepal CA-630 (NP-40), 0.5% sodium deoxycholate, 0.1% SDS, 1X protease inhibitor cocktail (complete Mini EDTA-free, Roche)). Lysates were sonicated (Bioruptor, Diagenode) and then quantified using Pierce BCA assay (Thermo Fisher). Identical amounts of proteins (20-40μg) were electrophoresed on 4–15% Mini-PROTEAN TGX precast protein gels (Bio-Rad). For transfer, the Trans-Blot Turbo transfer system (Bio-Rad) with nitrocellulose membranes was used according to the manufacturer’s instructions. TBS (137mM NaCl, 20mM Tris, pH 7.6) +5% nonfat milk was used for blocking, and TBS+0.1%Tween-20+5% milk was used for antibody incubations. An Odyssey infrared imaging system (LI-COR) was used to visualize blots.

### Flow Cytometry

Processed cells were flow cytometry analyzed immediately using either a BD FACSymphony A5 or BD LSRFortessa X-50 machine. Results were analyzed using FlowJo software.

### Viability Assays

Viable cell numbers were measured using CellTiter-Glo Luminescent Cell Viability Assay (Promega) according to manufacturer’s instructions.

### Xenograft tumors

All *in vivo* experiments were conducted in accordance with the NIH Guide for the Care and Use of Experimental Animals and with approval from the Fred Hutchinson Cancer Research Center, Institutional Animal Care and Use Committee (Protocol 1457). 100,000 GSCs were orthotopically xenografted into a single frontal cerebral hemisphere or in flanks of HSD:athymic nude Foxn1nu mice (#069, Envigo).

### Brain tumor gene expression analysis

Brain tumor and normal gene expression data were obtained from Children’s Brain Tumor Network [89], CGGA [90], GTEX [91], and TCGA [92]. A detailed description of creation of the analysis pipeline is available in (Arora et al. in preparation).

## Data availability statement

The RNA-seq data that support the findings of this study are openly available in the NCBI Gene Expression Omnibus at https://www.ncbi.nlm.nih.gov/geo/, GEO accession number GSE213269 (token: cfqhwiemrfifrqf). Other data that support the findings of this study are available in the Supporting Materials of this article.

## Ethical statement

We followed the guidelines set by the Fred Hutchinson Cancer Center Institutional Review Office for De-identified Human Specimens and/or Data, which categorizes the studies presented here as Research Not Involving Human Subjects as detailed by the Institutional Review Board’s Human Subjects Research Determination Form.

