## Supplemental Inventory and Figures for "Functional genomic analysis of adult and pediatric brain tumor isolates"

**Supplementary Information**

**Figure S1:** Principal component analysis of comprehensive GBM retest screens in human brain tumor isolates from Figure 1.

**Figure S2:** Gene set enrichment analysis for network hits and their neighboring nodes, in support of Figure 2.

**Figure S3:** Predictive feature expression analysis across diverse brain tumor types and also normal brain tissue in support of Figure 3.

**Figure S4:** FGF2 expression from in brain tumors, supporting Figure 4.

**Figure S5:** Data in support of Figure 6.

**Table S1**: Screen results from human brain tumor isolates

**Table S2:** RNA-seq data for screen isolates

**Table S3:** Functional genomic network integration of screen results

**Table S4**: Tumor key and expression of genes among brain tumors

**Table S5**: Key Resources

### Figure S1

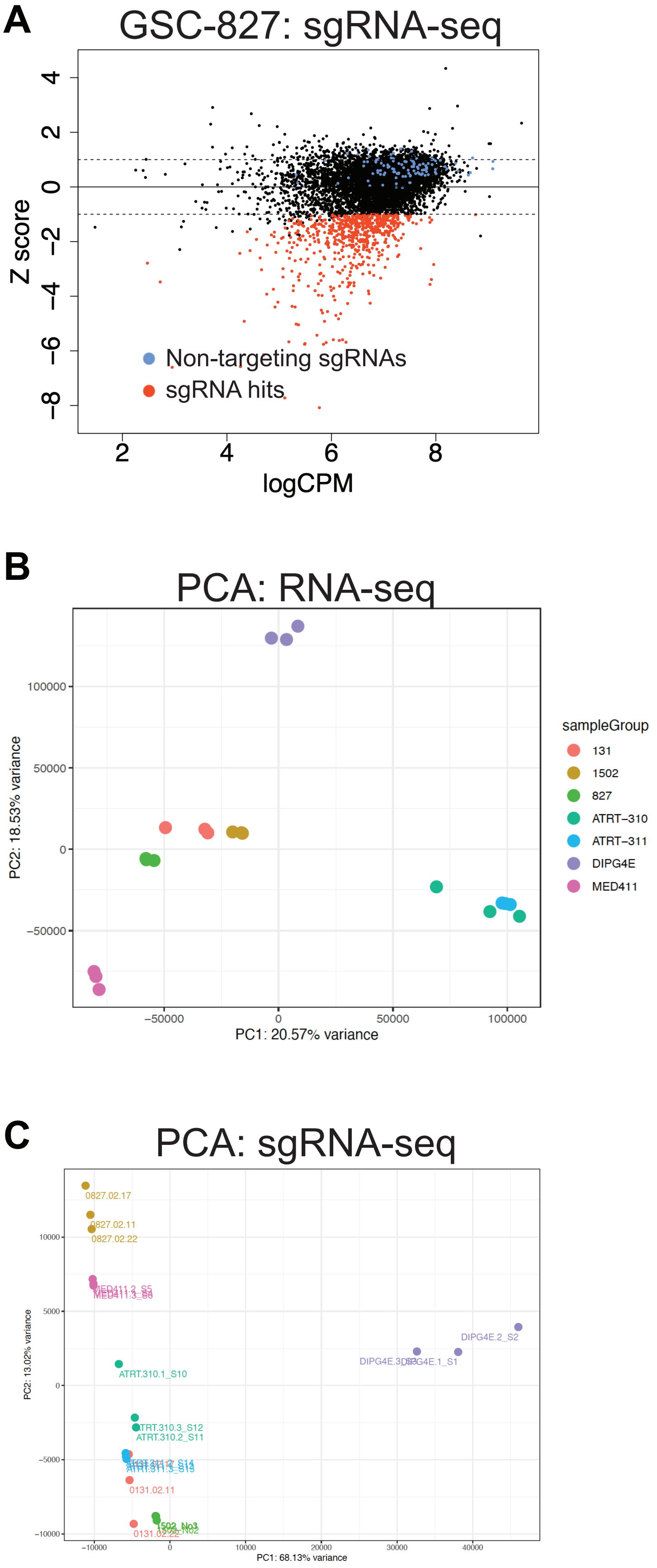


**Figure S1: Screen results and principal component analysis of comprehensive GBM retest screens in human brain tumor isolates (data supporting Figure 1).**
**(A)** sgRNA-seq analysis for GSC-0827 cells.
**(B)** RNA-seq data.
**(C)** sgRNA-seq data.

### Figure S2

**A**

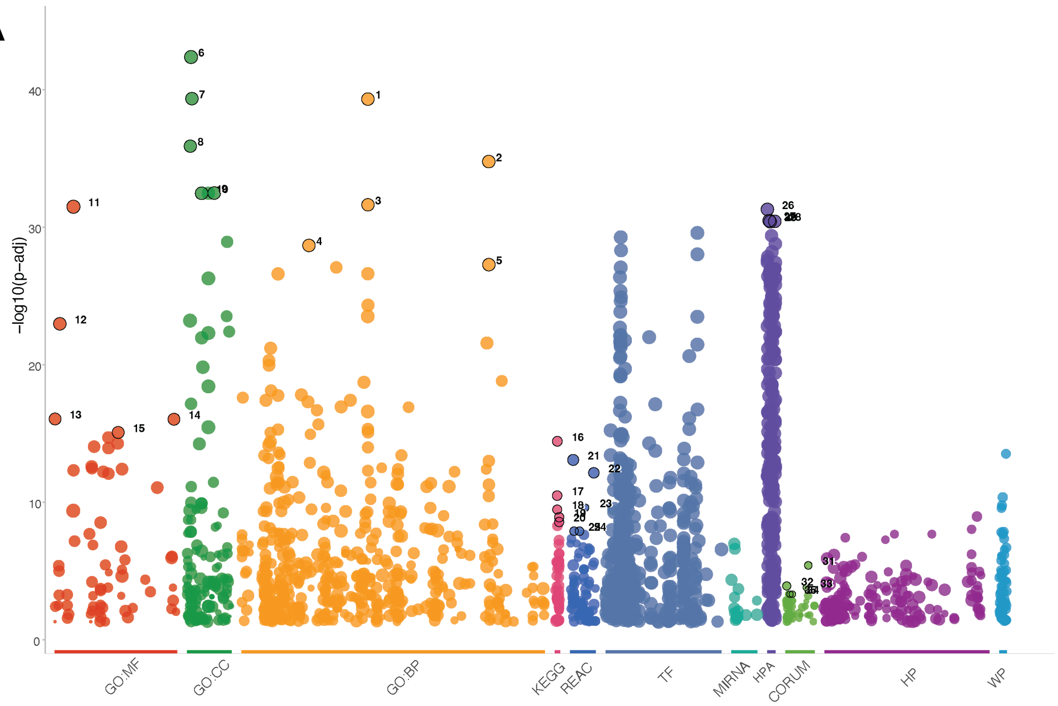


**B**

| id | source | term_id | term_name | term_size | p_value |
|---|---|---|---|---|---|
| 1 | GO:BP | GO:0044260 | cellular macromolecule metabolic process | 5766 | 4.8e−40 |
| 2 | GO:BP | GO:1901564 | organonitrogen compound metabolic process | 6446 | 1.7e−35 |
| 3 | GO:BP | GO:0044267 | cellular protein metabolic process | 4882 | 2.3e−32 |
| 4 | GO:BP | GO:0019538 | protein metabolic process | 5487 | 2.1e−29 |
| 5 | GO:BP | GO:1901576 | organic substance biosynthetic process | 5774 | 5.3e−28 |
| 6 | GO:CC | GO:0005737 | cytoplasm | 12241 | 4.2e−43 |
| 7 | GO:CC | GO:0005829 | cytosol | 5420 | 4.5e−40 |
| 8 | GO:CC | GO:0005654 | nucleoplasm | 4215 | 1.3e−36 |
| 9 | GO:CC | GO:0070013 | intracellular organelle lumen | 6625 | 3.3e−33 |
| 10 | GO:CC | GO:0031974 | membrane−enclosed lumen | 6626 | 3.4e−33 |
| 11 | GO:MF | GO:0005515 | protein binding | 14833 | 3.2e−32 |
| 12 | GO:MF | GO:0003824 | catalytic activity | 5746 | 1.1e−23 |
| 13 | GO:MF | GO:0000166 | nucleotide binding | 2176 | 8.7e−17 |
| 14 | GO:MF | GO:1901265 | nucleoside phosphate binding | 2177 | 9.2e−17 |
| 15 | GO:MF | GO:0036094 | small molecule binding | 2519 | 8.4e−16 |
| 16 | KEGG | KEGG:04150 | mTOR signaling pathway | 151 | 3.7e−15 |
| 17 | KEGG | KEGG:04110 | Cell cycle | 120 | 3.3e−11 |
| 18 | KEGG | KEGG:04115 | p53 signaling pathway | 72 | 3.4e−10 |
| 19 | KEGG | KEGG:05222 | Small cell lung cancer | 90 | 1.2e−09 |
| 20 | KEGG | KEGG:05215 | Prostate cancer | 93 | 2.7e−09 |
| 21 | REAC | REAC:R−HSA−1640170 | Cell Cycle | 678 | 8.3e−14 |
| 22 | REAC | REAC:R−HSA−3700989 | Transcriptional Regulation by TP53 | 362 | 7.2e−13 |
| 23 | REAC | REAC:R−HSA−8956320 | Nucleotide biosynthesis | 15 | 2.5e−10 |
| 24 | REAC | REAC:R−HSA−69236 | G1 Phase | 46 | 1.3e−08 |
| 25 | REAC | REAC:R−HSA−69231 | Cyclin D associated events in G1 | 46 | 1.3e−08 |
| 26 | HPA | HPA:0020052 | adrenal gland; glandular cells[...Medium] | 5981 | 5.0e−32 |
| 27 | HPA | HPA:0630000 | cervix | 6897 | 3.3e−31 |
| 28 | HPA | HPA:0550052 | stomach 2; glandular cells[...Medium] | 6252 | 3.8e−31 |
| 29 | HPA | HPA:0190221 | esophagus; squamous epithelial cells[...Low] | 6649 | 4.0e−31 |
| 30 | HPA | HPA:0190000 | esophagus | 6649 | 4.0e−31 |
| 31 | CORUM | CORUM:6664 | STAGA complex, SPT3−linked | 19 | 3.9e−06 |
| 32 | CORUM | CORUM:230 | Mediator complex | 32 | 1.2e−04 |
| 33 | CORUM | CORUM:6332 | Elongator complex (ELP1, ELP2, ELP3, ELP4, ELP5, ELP6) | 6 | 1.6e−04 |
| 34 | CORUM | CORUM:1237 | BAF complex | 9 | 4.8e−04 |
| 35 | CORUM | CORUM:807 | BRG1−associated complex | 9 | 4.8e−04 |

**Figure S2: Gene set enrichment analysis for network hits and their neighboring nodes, in support of Figure 2.**
**(A)** Graph of gprofiler analysis on network hits and their neighbors (Top5 terms for each category, using default parameters).
**(B)** Table showing specific categories.

### Figure S3

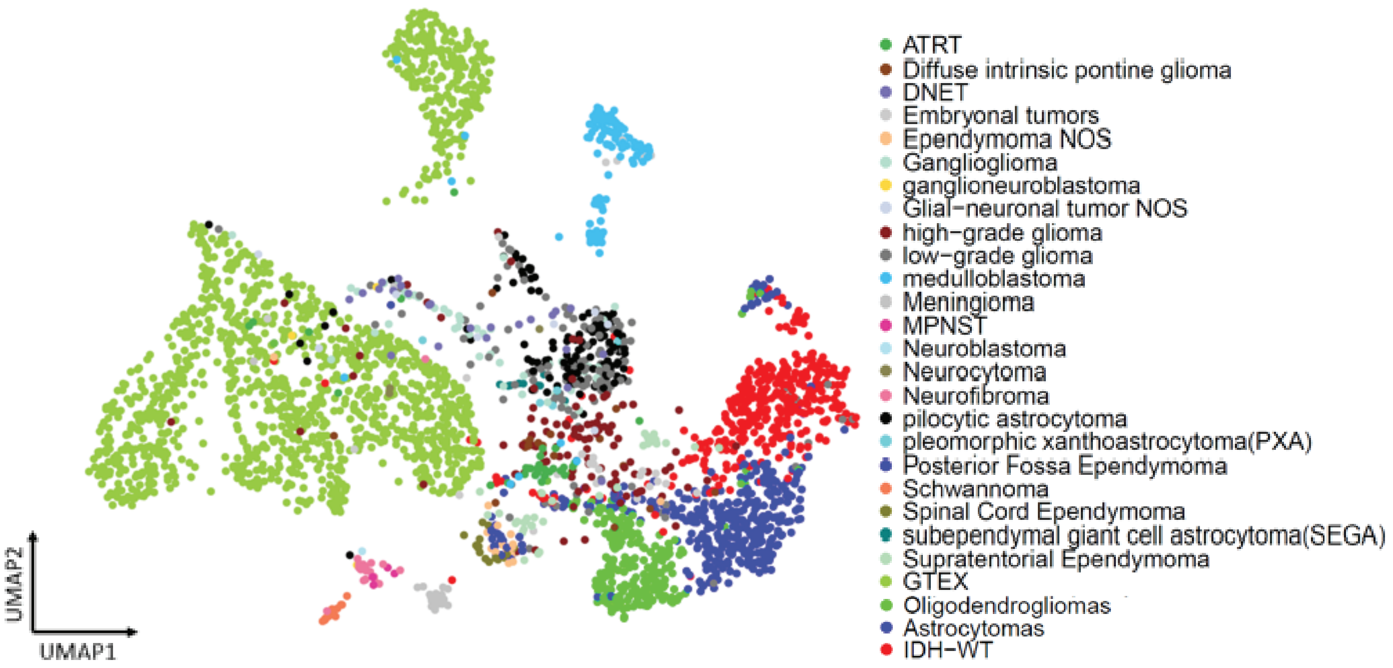


**Figure S3: Predictive feature expression analysis across diverse brain tumor types and also normal brain tissue.** UMAP projections of tumor and brain tissue samples used for gene expression analysis colored by tumors and tissue type.

### Figure S4

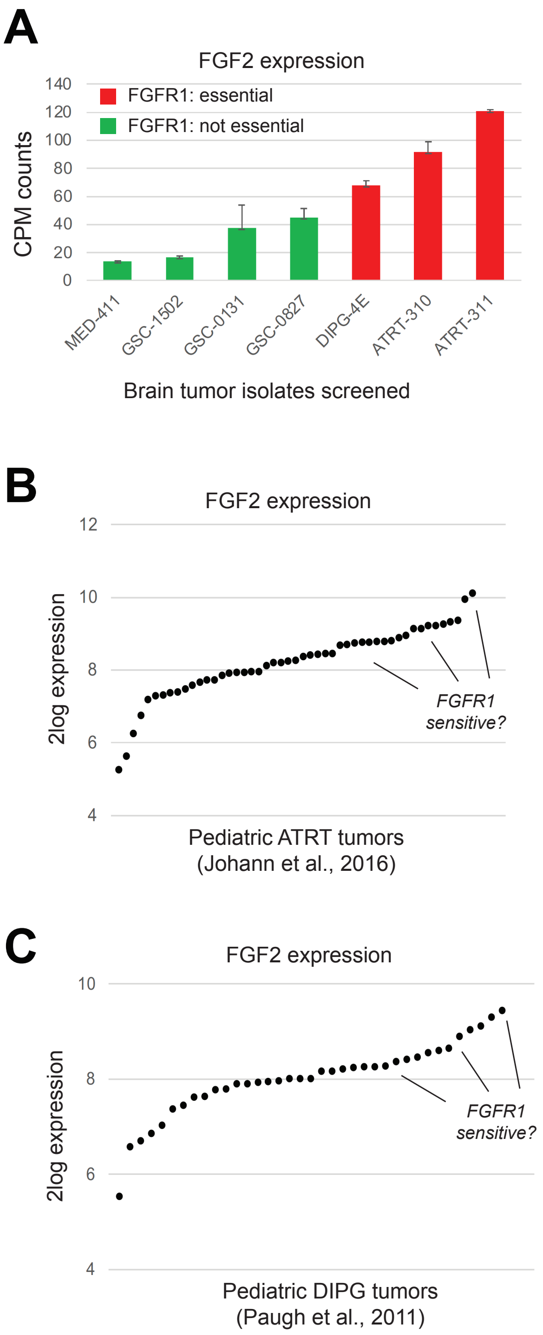


**Figure S4: FGF2 expression from in brain tumors, supporting Figure 4.**
**(A)** brain tumor isolates screened (RNA-seq).
**(B)** ATRT tumors (Johann et al., 2016; PMID:26923874).
**(C)** DIPG tumors (Paugh et al., 2011; PMID: 21931021).

**Figure S5: Data in support of Figure 6.**
**(A)** Heatmap of gene expression of genes associated with interferon responses patient-derived orthotopic xenograft (PDOX) models of pediatric brain tumors (Brabetz et al., 2018;PMID: 30349086).
**(B)** UMAP projections of tumor and brain tissue samples showing expression of MX1. Sample key is shown in Figure S4.
**(C)** Western blot analysis of ADAR, OAS1, and IRF9 for brain tumor isolates shown.
**(D)** Western blot analysis of MX1 in subset of pediatric brain tumor isolates and also **(E)** matched primary and recurrent adult GBM tumors. The results show that recurrent tumors can show significant MX1 expression, consistent with proneural to mesenchymal transition after SOC.

### Figure S5

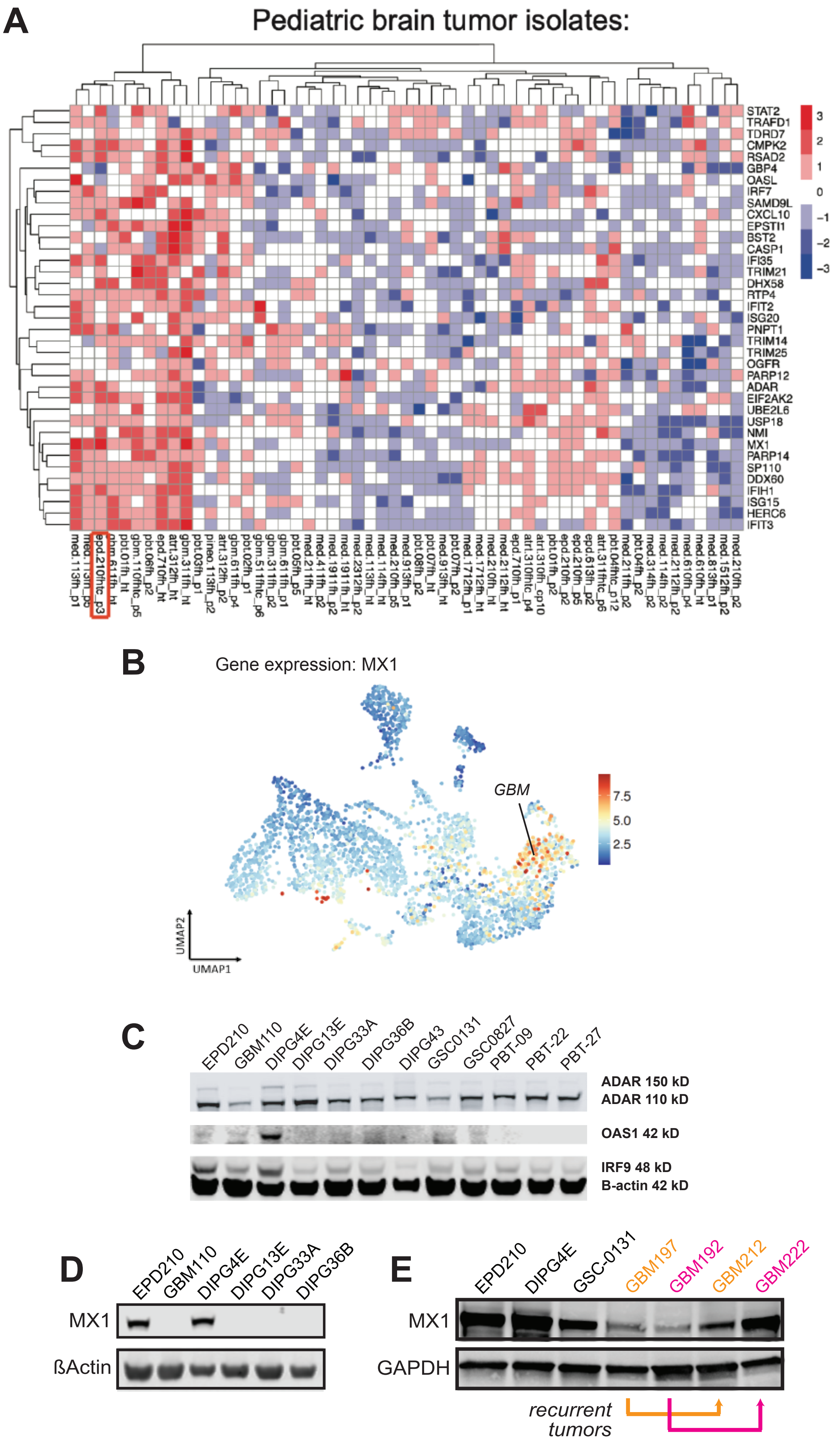
A
Pediatric brain tumor isolates:
STAT2
TRAFD1
TDRD7
CMPK2
RSAD2
GBP4
OASL
IRF7
SAMD9L
CXCL10
EPSTI1
BST2
CASP1
IFI35
TRIM21
DHX58
RTP4
IFIT2
ISG20
PNPT1
TRIM14
TRIM25
OGFR
PARP12
ADAR
EIF2AK2
UBE2L6
USP18
NMI
MX1
PARP14
SP110
DDX60
IFIH1
ISG15
HERC6
IFIT3
3
2
1
0
−1
−2
−3
B
Gene expression: MX1
GBM
7.5
5.0
2.5
UMAP2
UMAP1
C
EPD210
GBM110
DIPG4E
DIPG13E
DIPG33A
DIPG36B
DIPG43
GSC0131
GSC0827
PBT-09
PBT-22
PBT-27
ADAR 150 kD
ADAR 110 kD
OAS1 42 kD
IRF9 48 kD
B-actin 42 kD
D
EPD210
GBM110
DIPG4E
DIPG13E
DIPG33A
DIPG36B
MX1
ßActin
E
EPD210
DIPG4E
GSC-0131
GBM197
GBM192
GBM212
GBM222
MX1
GAPDH
recurrent
tumors
